# High accuracy methylation identification tools on single molecular level for PacBio HiFi data

**DOI:** 10.1101/2024.08.14.607879

**Authors:** Ying Chen, Bo Wu, Yu-Ying Ding, Long-Jian Niu, Xin Bai, Zhuo-Bin Lin, Chuan-Le Xiao

**Author notes:** To whom correspondence should be addressed: Chuan-Le Xiao. These authors contributed equally to the manuscript as first authors (Ying Chen, Bo Wu, Yu-Ying Ding, Long-Jian Niu).

## Abstract

PacBio’s Circular Consensus Sequencing (CCS) allows us to obtain highly accurate bases and simultaneously determine the methylation states of individual molecules. However, existing CCS-based methods for 5mC detection have low accuracy (<90% on most datasets) at the single-molecule level and can produce inaccurate methylation patterns. These methods rely on the information from 21 bp contexts surrounding the target CpGs and have over 29% low-confidence (<75% accuracy) calls at CpGs with less distinguishable signals. We hypothesize that incorporating CpG methylation correlation information at the single-molecule level could improve the methylation calls on low-confidence CpGs. Here, we present a novel deep graph convolutional network (hifimeth) that uses 400 bp context in CCS-based 5mC calling and show that its improved performance is mainly due to the inclusion of more neighboring CpGs in contexts. Hifimeth achieves an average single-molecule accuracy of 94.7% and an average F1 score of 94.2%, 5.5% and 5.9% higher than the previous state-of-the-art method, respectively. Hifimeth-based methylation frequency quantification by read counting outperforms previous methods on all human and zebrafish datasets tested. The results also show that hifimeth’s high-accuracy calls can reveal complex single-molecule methylation patterns, either related to haplotypes or repeat regions, with up to single-motif resolution.

## Introduction

DNA methylation is an epigenetic mechanism that alters chromatin structure and function without altering DNA sequence. It also interacts with other epigenetic features such as nucleosome positioning, histone modifications, and 3D genome organization (*1*). In addition, DNA methylation influences alternative splicing events, and distinct single-molecule methylation patterns can be associated with gene isoforms (*2*). To understand the complex epigenetic regulatory processes, it is essential to study DNA methylation patterns in chromatin at the single-molecule level, which represent the simultaneous methylation states of neighboring bases. In animals, DNA methylation occurs predominantly on cytosine bases, generating 5-Methylcytosines (5mCs) at CpG sites (*3*). 5mC methylation is critical for normal development and is involved in genomic imprinting (*4*), X-chromosome inactivation (*5*), transposable element silencing (*6*), and ageing (*7*). To elucidate the role of DNA methylation in these processes, it is necessary to accurately identify different methylation patterns between alleles and across repeated genomic regions. In the future, combining accurate single-molecule methylation detection with single-cell technology may help to reveal the detailed roles of DNA methylation in cell differentiation and carcinogenesis. Therefore, researchers have developed various technologies to faithfully unravel single-molecule methylation states (*8–10*).

Resolving the methylation states of individual molecules is a key challenge in DNA methylation research. Next-generation sequencing (NGS) methods such as bisulfite sequencing (BS-seq) are limited in their ability to capture single-molecule patterns due to their short read lengths. In contrast, single-molecule (long-read) sequencing technologies such as Pacbio and Nanopore can generate much longer reads and methylation-related signals from native DNA molecules without the need for additional DNA treatments (*10, 11*).

Nanopore sequencing can produce ultra-long reads of up to several megabases and has high a sensitivity of detecting 5mC (*12*). However, its high sequencing error rates can lead to loss of CpG sites and distortion of methylation patterns. The Circular Consensus Sequencing (CCS) mode of the recently released Pacbio Sequel II platform can generate high accuracy (>99%) reads with up to over 20 kb length by sequencing a single-molecule multiple times, which also helps to overcome the low signal strength of 5mC. Therefore, CCS is a promising technology for accurately revealing single-molecule methylation patterns. However, current CCS-based algorithms for 5mC detection at the single-molecule level have limited accuracy (<90% on most datasets tested) (*8, 13*). These algorithms infer the methylation states of CpG motifs from the polymerase kinetics information, which includes pulse width (PW) and inter-pulse duration (IPD). In addition, the kinetic signals are influenced by the sequence contexts (*14*), resulting in high signal complexity. Previous studies have shown that the kinetic information of the incorporated nucleotide is mostly correlated with the upstream 7 bases and the downstream 2 bases (*15*). Therefore, current algorithms have used short k-mer (≤ 21 bp) context information for methylation calling (*8, 13–16*). Among them, the deep learning-based method ccsmeth has been the most effective, almost reaching ∼0.90 accuracy (*8*).

Additional information may be required to further improve CCS-based CpG methylation detection. Previous studies have shown that neighboring CpG sites have significantly correlated methylation frequencies in mammals and plants (*17–19*). Based on this observation, some NGS-(*18, 20*) and long-read (*8*) based DNA methylation techniques have successfully improved the whole-genome site methylation frequency quantification. Alternatively, the correlation of methylation frequencies between neighboring CpGs could indicate the association of CpG methylation states at the single-molecule level, which could potentially assist methylation state calling. Therefore, we explored the single-molecule CpG methylation state association and developed a new deep learning-based method called hifimeth. For the first time, hifimeth leverages short range (< 250 bp) CpG methylation state correlation to improve CCS-based 5mC calling at the single-molecule level. On native human and zebrafish CCS data, hifimeth achieved an average accuracy of 94.7% accuracy and an average F1 score of 94.2%, an improvement of 5.5% and 5.9%, respectively, over the previous state-of-the-art method (ccsmeth) for 5mC detection at the single-molecule level; hifimeth-based whole-genome methylation frequency quantification through direct read counting was superior on all tested datasets, regardless of data source, sequencing library length, or sequencing depth. Furthermore, our analysis shows that the highly accurate CpG methylation calls enable the discovery of sophisticated single-molecule CpG methylation patterns, either allele-related or not.

## Results

### A potential improvement for methylation detection methods based on CCS

Pacbio’s CCS system sequences individual DNA molecules in multiple passes, potentially allowing methylation states of CpGs to be resolved with high accuracy. However, current CpG methylation calling algorithms use information from short k-mer (≤21 bp) contexts and have limited accuracy (<90% for most datasets), which can lead to biased single-molecule methylation patterns (Figure 1A,B). To improve methylation calling, we evaluated the limitations of current methods. Primrose and ccsmeth estimate methylation probabilities for each CpG with kinetic signals. We evaluated the accuracy of their positive and negative predictions (predictive values) at different probability thresholds (Figure 1B). For both algorithms, applying stricter separate thresholds for positive and negative calls increased the positive and negative predictive values but reduced data utilization (Figure 1B). For example, on a Pacbio Sequel II flowcell of the human cell line HG002, changing the threshold from the default 0.5 to ≤0.10 for negative calls and ≥0.90 for positive calls increased the positive / negative predictive values from 0.790 / 0.898 (primrose) and 0.889 / 0.927 (ccsmeth) to 0.924 / 0.970 (primrose) and 0.978 / 0.984 (ccsmeth), at the expense of losing 40.9% and 29.6% read CpG sites, respectively. A similar trend was observed for the K-20 model of hifimeth (see Methods and Supplementary Figure 1A). Consequently, we considered the CpG methylation calls with ≤0.10 or ≥0.90 methylation probabilities as high-confidence, while the rest with an overall accuracy of 0.706 (primrose) and 0.745 (ccsmeth) as low-confidence (Supplementary Figure 1B). These results suggest that the low single-molecule accuracy of the short k-mer models is mainly due to the low-confidence CpG calls.

**Figure 1.**
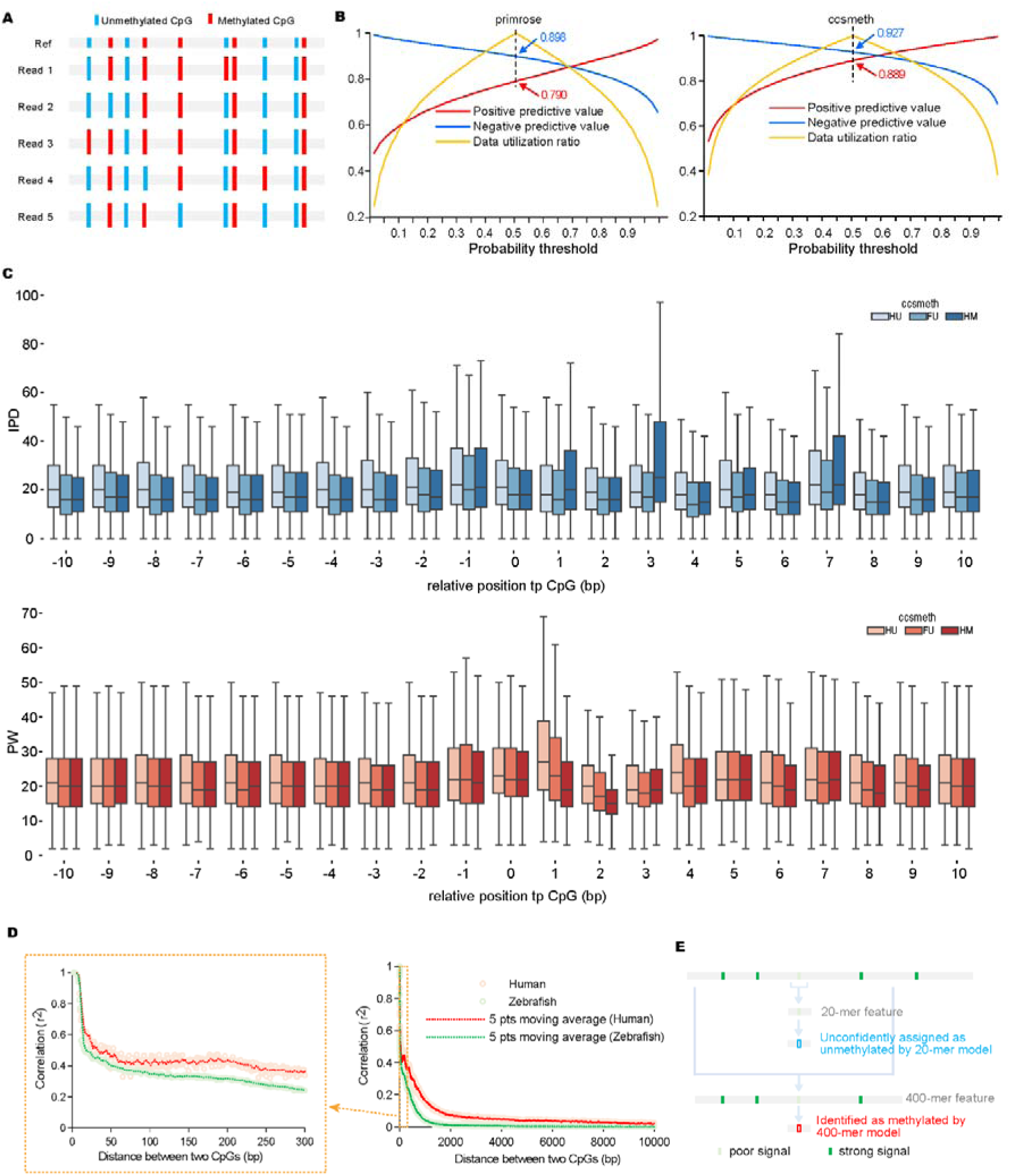
Current challenges and a possible solution for CCS-based methylation detection methods. (**A**) Schematic diagram illustrating how ∼90% methylation detection accuracy affects the observed single-molecule patterns. (**B**) The performance of primrose (left) and ccsmeth (right) using different software methylation probabilities as thresholds. Positive (negative) predictive value is the proportion of true positive (negative) calls among all positive (negative) calls. Positive and negative predictive values are calculated using a single probability threshold, with probabilities ≥ and < the threshold classified as positive and negative calls, respectively. The red / blue arrows indicate the positive / negative predictive values at the 0.5 threshold. The data utilization ratios are calculated using symmetric thresholds for positive and negative calls; when α (< 0.5) is the negative threshold, (1 – α) will be the positive threshold, ≤ α probabilities are classified as negative, and≥ (1 – α) probabilities are classified as positive. Accordingly, for data utilization ratio, the points α and (1 - α) correspond to the same threshold pair on the horizontal axis. (**C**) Distribution of inter-pulse duration (IPD) and pulse width (PW) signals in 21 bp contexts of ccsmeth high and low confidence CpGs. HU, correct high confidence unmethylated CpGs; HM, correct high confidence methylated CpGs; FU, false negative low confidence CpGs. (**D**) Correlations of methylation states among CpGs at the single molecule level within 300 bp (left) and 10000 bp (right) distances. (**E**) A potential strategy that uses methylation state correlation to improve CCS-based single-molecule methylation calling.

Further improvements depend on more accurate calling of the low-confidence CpGs. To explore this possibility, we compared the 21bp context kinetic signals (IPD and PW) of correctly called methylated and unmethylated high-confidence CpGs with those of the false negative low-confidence CpGs by ccsmeth. As shown in Figure 1C, at most upstream positions (-10 to -2), the motif base C and a few downstream positions (4, 8, 9, and 10), the IPD and PW signals for the false unmethylated CpGs are more similar to the methylated high-confidence calls, while at the remaining positions their signals are either closer to the unmethylated CpGs or similarly distant to the methylated and unmethylated CpGs. These results suggest that the false negative CpGs have distinct context signals from the unmethylated CpGs, but not as clear as the high-confidence methylated CpGs.

The association of methylation states between CpGs on single molecules can provide additional information for determining the methylation states of low-confidence CpGs. In mammalian genomes, CpG sites tend to cluster and form CpG-rich regions such as CpG islands (*21*), which tend to have synchronized methylation states among internal CpGs. Previous studies have also found a significant correlation of methylation frequencies between neighboring CpG sites in both animals and plants (*22*). Based on the high-confidence CpG calls, we confirmed that the methylation states of adjacent CpGs sites are significantly correlated in single molecules in both human and zebrafish (Figure 1D and Supplementary Note 1). Therefore, we hypothesize that incorporating the methylation (kinetics) information of adjacent CpGs could improve methylation calls on low-confidence CpGs (Figure 1E).

### Design of hifimeth

Here, we developed hifimeth to detect 5mC with high accuracy at the single-molecule level by incorporating more neighboring CpGs to support methylation calling (Figure 2A). We first considered using short contexts of both the target CpG and a certain number of neighboring CpGs. However, this strategy was abandoned mainly due to the large variance of CpG density in the human genome (Figure 2B), which renders neighboring CpGs useless in CpG sparse regions and the contexts overlap in CpG dense regions. For instance, using 20 bp k-mers, the context overlap is estimated to be no less than 37.8% in the human genome (Figure 2C). Analyses in Supplementary Note 1 show that CpGs within less than 250 bp distances in human or zebrafish have strong correlation (*r^2^* ≥ 0.40 and FDR<0.05) (Figure 1D). Therefore, we adopted a second strategy to extend the context of the studied CpG, whose length is positively correlated with the chance of including additional CpGs (Figure 2C). The first challenge in training a long context deep learning model is the significant increase in the computational load compared to the short context model. We ran out 256 GB of memory when trying to train a 400 bp k-mer model using ccsmeth, which would also take a long time (several days) even with sufficient memory (>1 TB). As a result, we redesigned deep graph convolutional (*23*) neural networks (hifimeth) for CCS-based 5mC detection (Figure 2A), which consumes much less memory and training time, taking only a few hours to train a 400 bp model.

**Figure 2.**
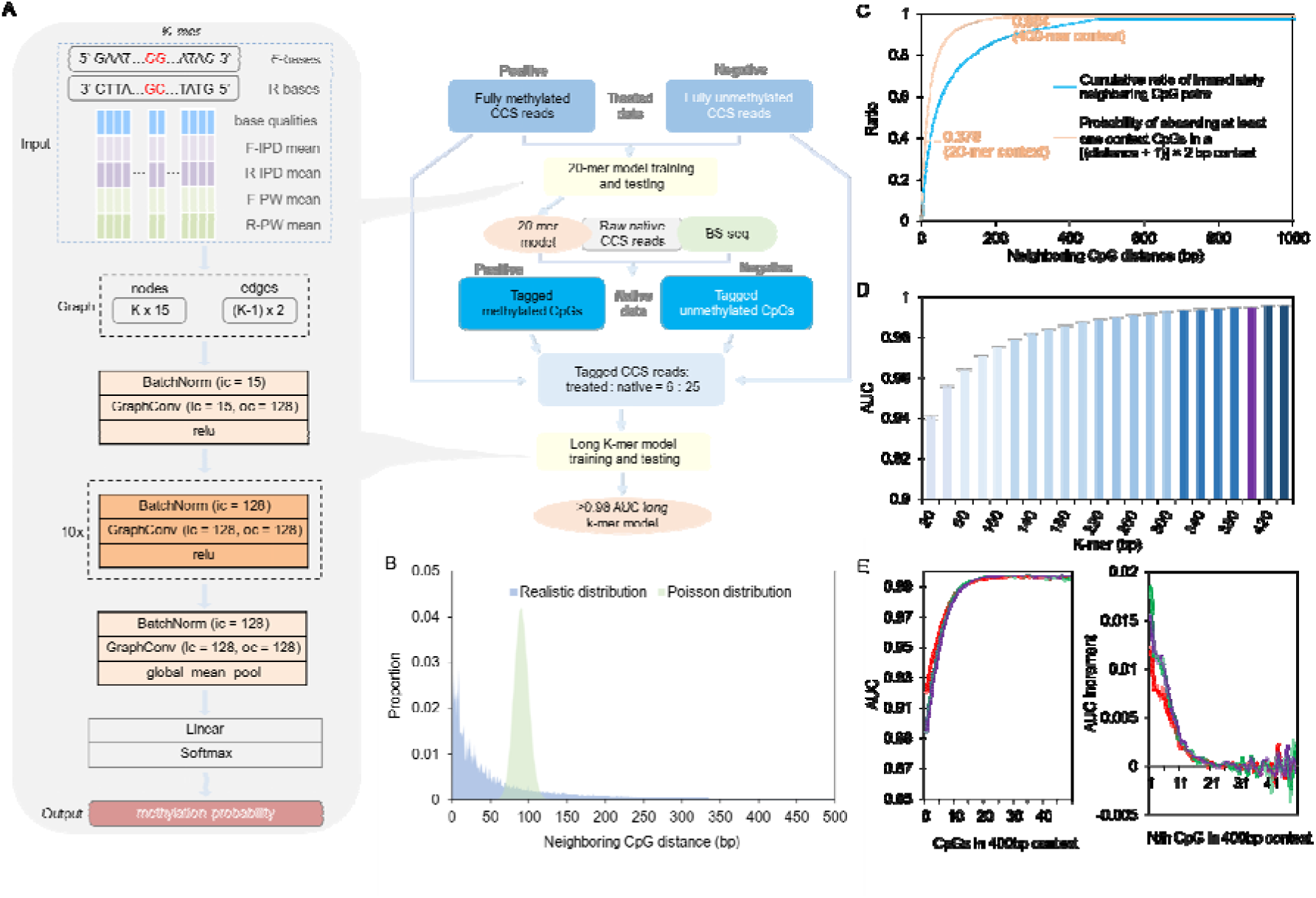
Effects of context length and CpG density on the performance of deep neural networks for DNA methylation inference. (**A**) Schematic diagram of the graph convolutional neural networks (left) and the training processes (right) of hifimeth. (**B**) Comparison of the realistic and simulated stochastic (Poisson distribution) CpG motif distribution in the human genome. (**C**) Cumulative ratios of distances between CpGs and their first upstream or downstream CpGs in the human genome (blue curve). The probability (orange curve) of observing at least one extra CpG at different context lengths [equaling to (distance + 1) × 2 in graph] is calculated based on the observed cumulative ratios (blue curve). (**D**) Areas under ROC curve (AUC) of hifimeth models using different context k-mer lengths. All models were trained and tested (three repeats) using the same datasets. (**E**) AUC of the 400 bp hifimeth model on CpGs with different numbers of extra CpGs (left) and AUC gain of each additional extra CpG (right) in context.

We then attempted to identify the optimized context length for 5mc detection by hifimeth. We trained hifimeth with different k-mers ranging from 20 bp to 440 bp with a 20 bp increase per step using a same native training dataset. The results show that the AUC (area under the ROC curve) score of the model increases with the k-mer length (Figure 2D), confirming that the applied context length has a major impact on the CpG methylation detection. The increase of AUC score slows down with increasing the k-mer length, approaching the maximum at approximately 400 bp. Considering the limited gain in benefit and the increase in computational burden of further increasing the context length, we selected 400 bp as the optimal context length for hifimeth and trained the k-400 model. The improved performance of hifimeth may be due to the incorporation of additional neighboring CpGs. To test this hypothesis, we evaluated the AUCs of the hifimeth k-400 model on CpGs with different numbers (0 to 49) of context CpGs. As shown in Figure 2, tested on six flowcells of native HG002 CCS datasets, the AUC of the hifimeth k-400 model increases from a minimum of 0.904 ± 0.012 (0 extra CpG in context) to a maximum of 0.996 ± 0.001 on CpGs with ≥21 extra context CpGs. The effect of context CpGs approaches saturation at approximately 21. And the AUC improvement efficiency of extra CpGs decreases rapidly from the 1^st^ (0.015 ± 0.002 AUC increase) to the 21^st^ extra CpG (<0.001 AUC increase) (Figure). In humans, there is a 98.4% probability of containing at least one extra CpG site in a 400-mer context (Supplementary Figure 1), so the k-400 model can theoretically can assist the methylation calling of most CpG sites.

To learn the knowledge of native CpG methylation association, we mainly used native human CCS reads to train the k-400 model (see Methods). As shown in the following sections, with all the above optimizations, hifimeth has achieved dramatically improved single-molecule level accuracy and methylation frequency quantification quality by read counting compared to previous methods.

### Hifimeth outperforms short k-mer models at the single-molecule level

To confirm the improved single-molecule performance of hifimeth, we compared it with two 21-mer based methods, primrose and ccsmeth, on native CCS datasets at the read level. The datasets tested included 9 flowcells of human CCS reads, and one flowcell of zebrafish CCS reads. The tests used whole-genome fully methylated and unmethylated CpGs sites in BS-seq (for human cell line HG002 and zebrafish) or nanopore (for human cell line CHM13) data as reference positive and negative CpG sites. For all datasets tested, hifimeth outperformed ccsmeth and primrose in terms of precision, accuracy, recall rate, and F1 score (Figure 3A). The AUCs by hifimeth (0.987 ± 0.004) are higher and more stable (lower variance) than primrose (0.913 ± 0.009) and ccsmeth (0.961 ± 0.010) on all datasets tested (Figure 3B and Supplementary Table 1). On the human data, hifimeth has an average accuracy of 94.6% ± 0.8% and an average F1 score of 94.2% ± 0.6%, higher than primrose (83.3% ± 1.0% accuracy and 81.3% ± 1.1% F1 score) and ccsmeth (89.5% ± 1.1% accuracy and 87.9% ± 0.9% F1 score). Although we trained the K-400 model on human data, it has higher accuracy (95.8%) and F1 score (97.5%), and a greater improvement in accuracy compared to primrose (82.7% accuracy) and ccsmeth (86.5% accuracy) on the flowcell of zebrafish CCS reads. The zebrafish genome (17.4 CpGs/kb) has ∼1.62 times the CpG density of the human genome (10.7 CpGs/kb in CHM13 v2.0), which may explain the improvements.

**Figure 3.**
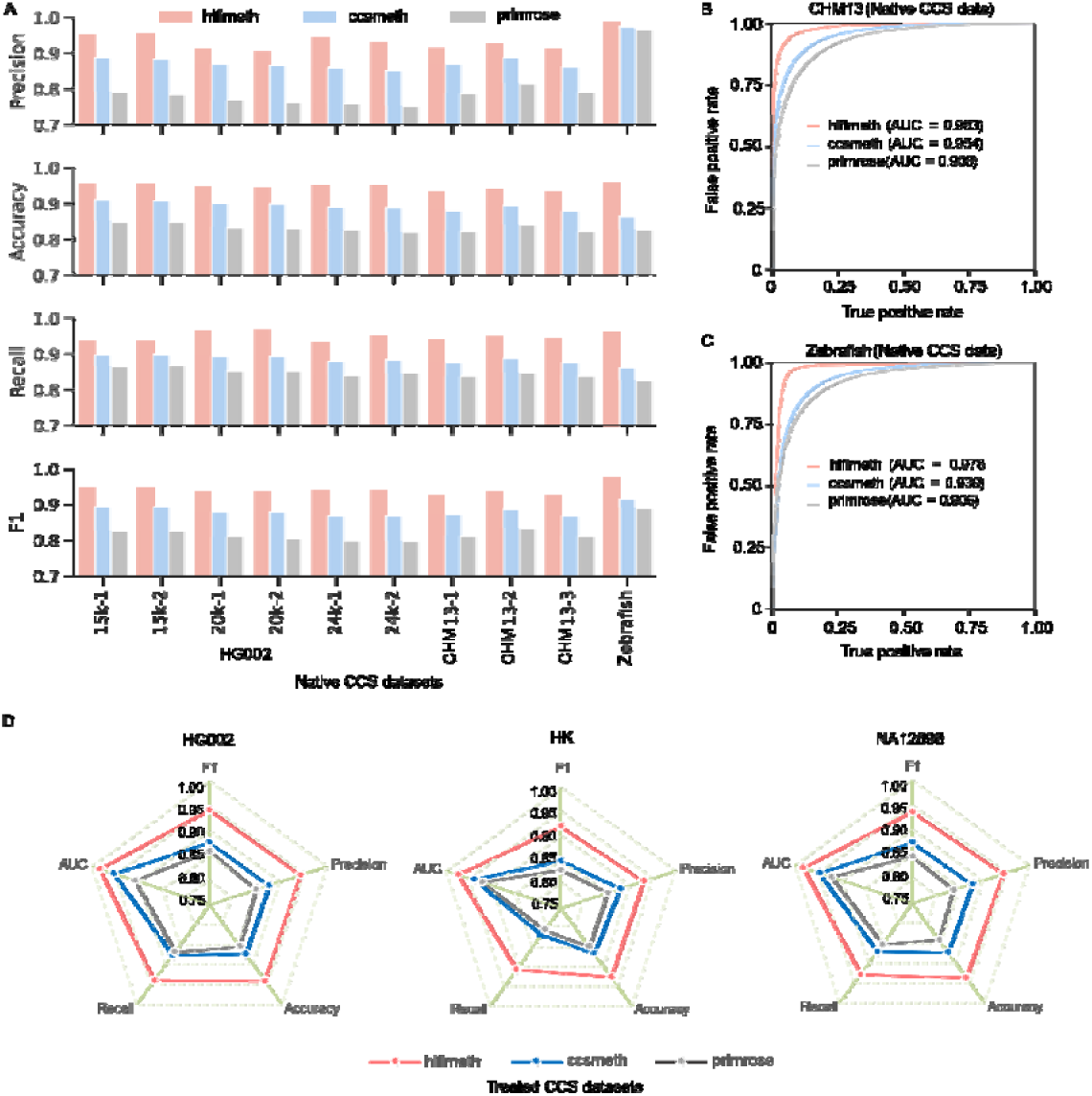
Performance comparison of hifimeth, ccsmeth and primrose at the single-molecule level. (**A**) Performance of the three softwares on native human and zebrafish CCS data. The graph includes assessment results on 1 flowcell of zebrafish data and 9 flowcells of human data, including 6 flowcells of HG002 cell line (2 per 15 kb, 20 kb, or 24 kb library size) and 3 flowcells of the CHM13 cell line. (**B** and **C**) ROC curves and areas under curves (AUC) of the three softwares on CHM13 and Zebrafish data. For CHM13, average scores of the three flowcells are shown in the graph. (**D**) Comparison of hifimeth, ccsmeth and primrose on three treated datasets. Treated datasets include CCS reads from DNAs fully methylated by treating with methyltransferase as positive control and those fully unmethylated via PCR amplification as negative control.

We also evaluated the strength of hifimeth at the read level using three CCS datasets from treated DNAs of different human cell lines, HG002 (partial training data of ccsmeth and hifimeth) (*8*), NA12898 (training data of primrose) (*8*), and M2&W2 from Tse. et al (*13*). These treated DNAs are composed of CpG fully methylated reads and fully unmethylated reads whose source DNAs were treated with M.SssI methyltransferase and PCR amplification, respectively. Despite being trained mainly on native CCS reads, the k-400 model of hifimeth performs better than primrose and ccsmeth which used larger proportions of treated data in training on all three tested datasets (Figure 3C and Supplementary Table 1). Overall, the hifimeth k-400 model has dramatically improved performance at the single-molecule level compared to primrose and ccsmeth, and its advantage is broadly present across different types of CCS data.

### High single-molecule-level accuracy improves CpG site methylation frequency quantification

An important purpose of methylation detection is to quantify the methylation frequencies across the genome. Higher single-molecule accuracy and recall rate will theoretically lead to better site frequency quantification results. To test this hypothesis, we sampled subsets of CCS reads at 10×, 15×, 20×, and 25× sequencing depths for four types of libraries, including HG002 15 kb, 20 kb (only sufficient for 10× and 15× sampling), 24 kb and CHM13 20kb. We then compared primrose, ccsmeth, and hifimeth in quantifying methylation frequencies of whole-genome CpG sites by read counting. We assessed their quality by correlating with results from ∼100× BS-seq for HG002 or ∼33× nanopore data for CHM13. The results showed that hifimeth had the highest correlation coefficients with either BS-seq or nanopore sequencing-based analysis across all tested datasets, followed sequentially by ccsmeth and primrose (Figure 4A). Hifimeth had greater advantages over ccsmeth and primrose on datasets at lower sequencing depths (Figure 4A). With 15× CCS reads, hifimeth had ≥ 0.90 correlations with either the BS-seq (HG002) or ONT (CHM13) quantification results on most (3/4) libraries tested. On the HG002 24kb and CHM13 20kb libraries, hifimeth with 10× or 15× reads had outperformed ccsmeth and primrose with 25× reads. We also compared their whole-genome site frequency quantification results by read counting on ∼23.3× zebrafish CCS data, on which hifimeth had a 0.075 and 0.110 correlation coefficient improvement compared to ccsmeth and hifimeth, respectively (Figure 4B).

**Figure 4.**
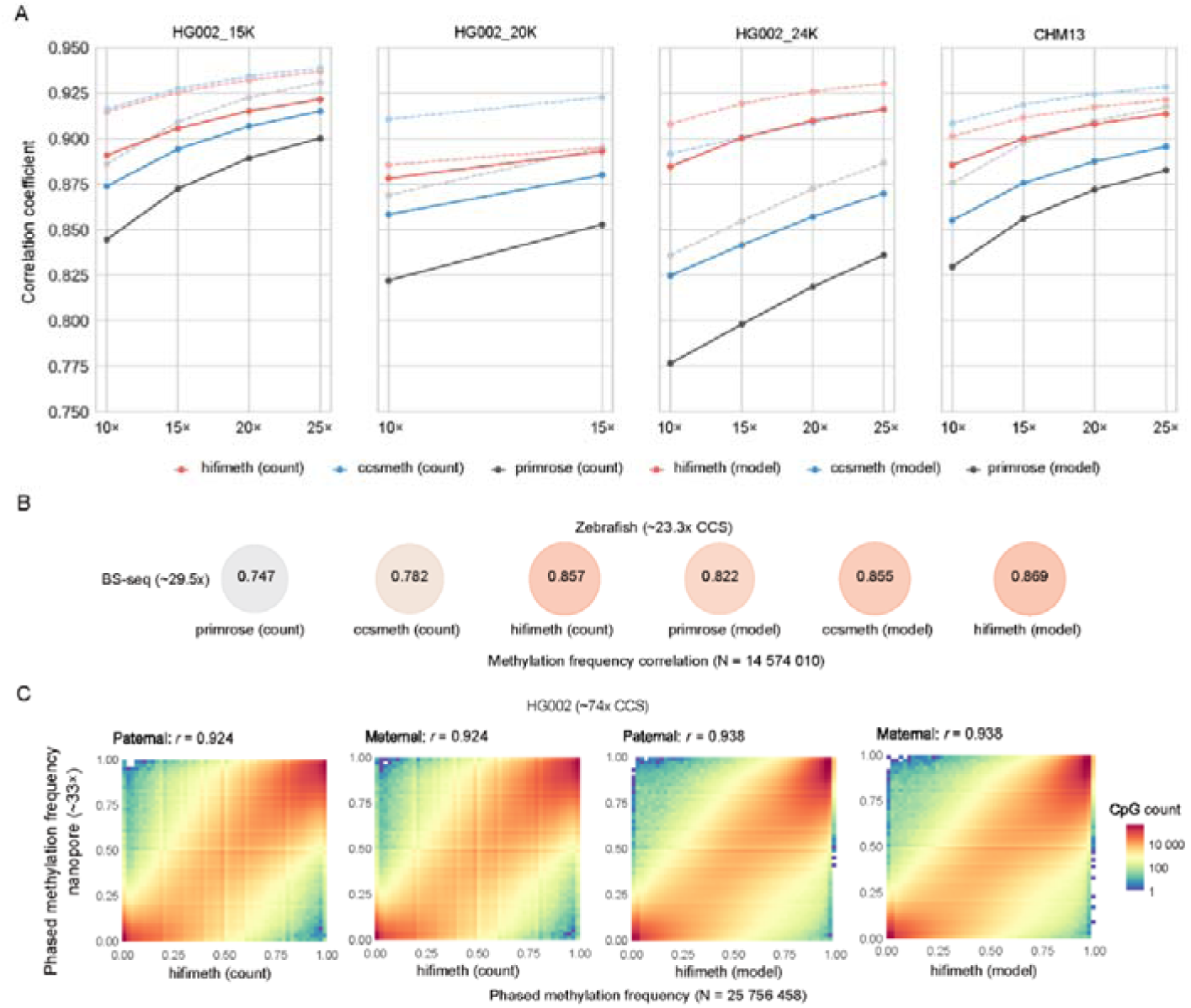
Whole-genome CpG methylation frequency quantification by hifimeth. In parentheses after software names, count and model denote quantification using the read counting and site frequency model modes, respectively. (**A**) Correlation coefficients (y-axis) of methylation frequency quantification by hifimeth, ccsmeth, and primrose, with reference quantification results of nanopore sequencing on human CCS datasets of varying sequencing depths (x-axis). The CCS data were randomly downsampled from three HG002 libraries with different insert sizes (15 kb, 20 kb, and 24 kb) and one CHM13 library (20 kb). For tests on < 25× depth data, the mean correlation coefficients of three repeats are shown, and individual test results can be found in Supplementary Table 2. (**B**) Comparison of whole-genome methylation frequencies obtained by six methods on a zebrafish CCS data flowcell with BS-seq data using correlation coefficients. (**C**) Heatmap and correlation analysis of phased (paternal and maternal haplotypes) methylation frequencies of HG002 between CCS-based hifimeth and nanopore-based analyses.

Ccsmeth and primrose have implemented site frequency models that incorporate the methylation frequencies of upstream and downstream CpG sites for better correlation with BS-seq data. Hifimeth’s read count model performed similarly to ccsmeth’s site-frequency model on the HG002 24kb library and the zebrafish data. Nevertheless, we trained a site-frequency model for hifimeth using the method used by ccsmeth (*8*), which resulted in slight improvements in correlation coefficients with BS-seq or nanopore results on all tested datasets (Figure 4A). The hifimeth frequency model mode outperformed primrose on all datasets tested, while tied with the ccsmeth model mode. The results of the hifimeth model mode were better than the ccsmeth model mode on two datasets (HG002 24kb library and the zebrafish data), similar on the HG002 15kb library and worse on the HG002 20kb library and the CHM13 library (Figure 4A,B).

Given phased variations or haplotype-tagged bam files, hifimeth can perform haplotype-level site methylation frequency quantification. We evaluated the haplotype-level quantification performance of hifimeth using HG002 CCS reads. From 10× to 74× sequencing depth, the correlation coefficient between the methylation frequencies using hifimeth count mode and those based on ∼40× nanopore data increased from 0.864 (10× depth) to 0.924 (74× depth) on either the paternal or maternal haplotype, and from 0.893 to 0.938 using the frequency model mode (Figure 4C and Supplementary Figure 2); the number of quantified CpG sites (≥5 reads per haplotype) also increased from 7.05 M (million) to 24.8 M (Supplementary Table 2). At ∼30× depth, the correlation coefficient reached 0.903 (count mode), and the quantified CpG sites exceeded 95% (23.6 million) of those quantified at ∼74× depth (Supplementary Figure 2 and Supplementary Table 2). Based on hifimeth calls, we detected 15 908 allelic differentially methylated regions (DMRs) with ≥20% allelic difference, 65.6% of which overlapped with nanopore-based allelic DMRs (Supplementary Table 3). And the distribution of hypermethylated paternal and maternal regions across the genome is significantly correlated (p<0.001, *r* = 0.855 and 0.874) between CCS-hifimeth and nanopore-based results (Figure 5A). These allelic DMRs also overlap with 79.7% (114) of 143 (Supplementary Table 4) well-supported imprinted DMRs reported by Akbari et al. (*24*).

**Figure 5.**
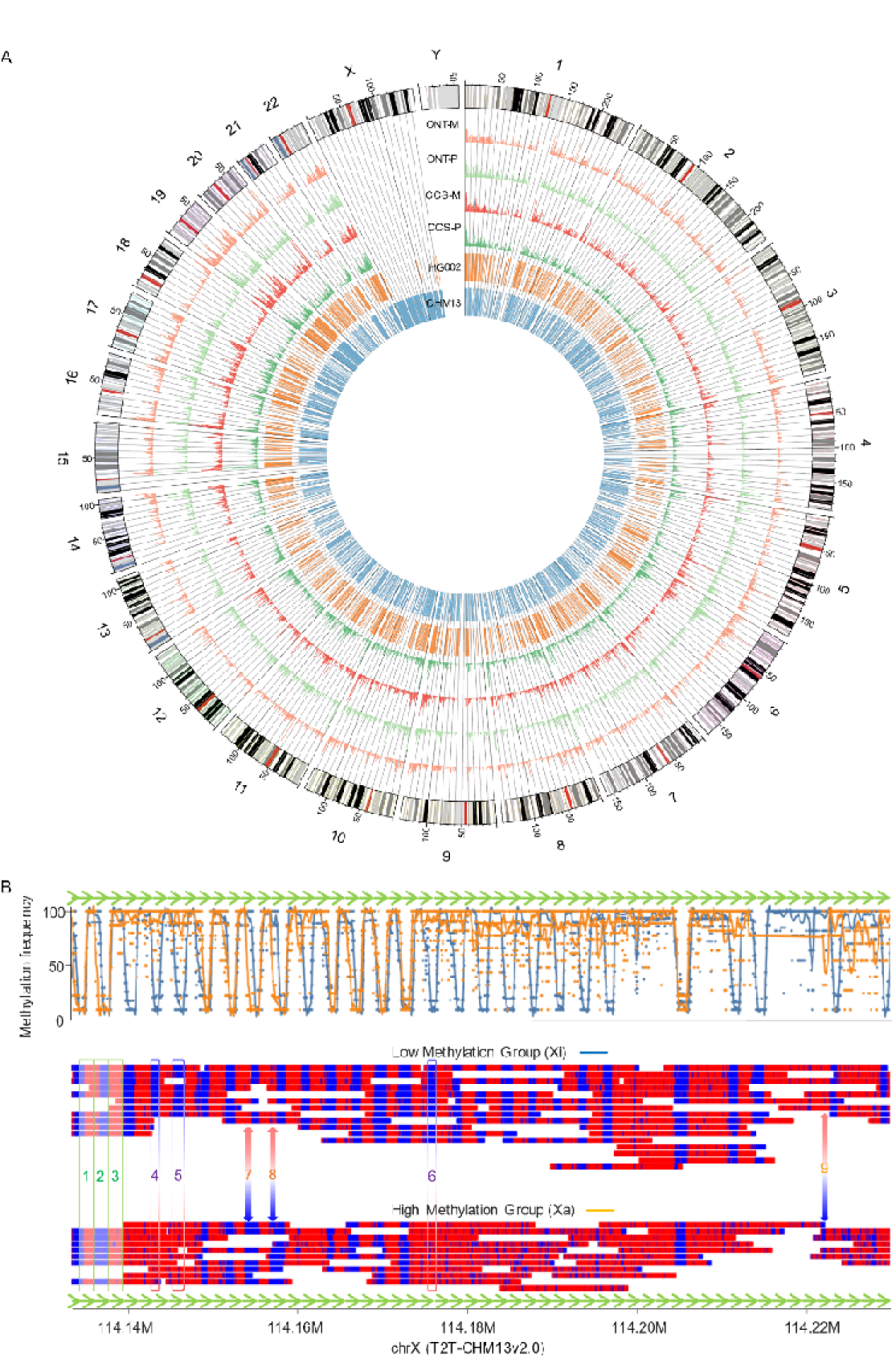
Differential methylation analysis by hifimeth. (A) Genome-wide distribution of allelic differentially methylated regions (DMRs) and read methylation-clustered regions. The outer ring shows the 24 chromosomes of GRCh38 with cytogenetic bands from UCSC genome browser. The next four histogram layers show the proportions of maternal (-M) or paternal (-P) hypermethylated regions detected by nanopore sequencing (ONT) and CCS with hifimeth in 1 Mb windows (vertical axes ranging from 0 to 0.01). The inner ring shows the read methylation-clustered regions (≥ 400 bp) in the HG002 (∼74×) and CHM13 (∼25×) CCS datasets. (B) Methylation patterns in the DXZ4 tandem repeat region of CHM13 by hifimeth based read clustering. Each green arrow represents a ∼3 kb repeat unit. The top panel shows the methylation frequencies in the low and high methylation groups on each CpG (dots), and the smoothed average methylation frequencies (curves) by modbamtools (Razaghi et al., 2022). The bottom single-molecule plots show the methylation states (red, methylated; blue, unmethylated) of CpGs in each read of the low and high methylation groups, inferred to be the inactivated X chromosome (Xi) and active X chromosome (Xa), respectively. In the bottom panels nine exemplary regions are marked: green frames (1–3): variable methylation across repeat units; blue-red frames and blue/red double arrows: regions with consistent (4–6) and opposite (7–9) clustering to the read group assignment, respectively.

Hifimeth calls the methylation status of all CpGs on CCS reads with kinetic information like ccsmeth, and has the same advantages in genome coverage over NGS-based methods, especially in repeat regions (*8*). Overall, hifimeth enables high-quality whole-genome methylation frequency quantification with relatively low CCS sequencing depth due to its improved performance at the single-molecule level.

### Unravelling complex single-molecule CpG methylation patterns with hifimeth

High-accuracy single-molecule methylation detection enables the exploration of diverse methylation patterns among homologous sequences. Methylation-based clustering can separate allelic reads in imprinted genomic regions without using phased genetic variants (*25*). In this study, we attempted to differentiate paternal and maternal reads in genomic regions exhibiting ≥40% allelic methylation solely based on methylation calls by hifimeth calls. To achieve this, we divided the reference genome into small (200 bp) bins and categorized the reads into high-methylation groups (HMGs) and low-methylation groups (LMGs) (see Methods). We considered bins or regions (merged clustered bins) with a majority (≥80%) of overlapped reads assigned to HMG or LMG as clustered bins or regions, respectively. On the HG002 CCS dataset (∼74×), we identified a total of 9 738 clustered bins and 1 259 clustered regions. Remarkably, 6 822 (70.0%) of the clustered bins and 1 179 (93.6%) of the clustered regions overlapped with allelic DMRs (Figure 5A and Supplementary Table 5), indicating that methylation-based clustering effectively captured allelic differences in methylation patterns. Furthermore, within the promoter regions of 727 protein encoding genes (Supplementary Table 6), we discovered at least one clustered bin, thereby identifying potential imprinted genes, including 24 previously reported ones (*24*). For instance, *CYREN,* undocumented as an imprinted gene before, harbored two clustered bins in its promoter and genic region in HG002, with paternal reads predominantly assigned to HMG and maternal reads assigned to LMG (Supplementary Figure 3). Similar HMG and LMG reads of *CYREN* were also observed in CHM13.

Methylation-based read clustering theoretically has the potential to distinguish reads in low-allelic-variance or homozygous genome regions. Out of the 80 clustered regions that do not overlap with any allelic differentially methylated regions (DMRs), 42 regions had fewer than 5 haplotype-tagged reads in at least one haplotype due to low local allelic variance (Supplementary Table 5). CHM13, a cell line with an almost homozygous genome and a female karyotype, still demonstrated the presence of genomic imprinting effects (26). Using hifimeth calls on approximately 25× CCS reads, we identified a total of 3,006 clustered regions (Supplementary Table 7), with 1,376 of them located on the heavily imprinted chromosome X, exhibiting an enrichment of approximately 15.6-fold compared to the autosomes (p < 0.001 by Fisher’s exact test). Within the DXZ4 tandem repeat region of chrX (Figure 5B), the low-methylation group (LMG) and high-methylation group (HMG) reads successfully captured the differential methylation patterns of the active and inactivated X chromosomes, similar to the observations made using approximately 33× nanopore reads (*26*). These results highlight the ability of hifimeth calls to effectively distinguish allelic methylation patterns in regions with low allelic variance levels.

By utilizing CCS-hifimeth calls, methylation-based read clustering can uncover differential methylation patterns that are not necessarily related to allelic differences. Among the remaining 38 clustered regions in HG002 that do not overlap with any allelic differentially methylated regions (Supplementary Table 5), 14 of them are located on chromosome X (Figure 5A). Since HG002 is a male cell line, these 14 clustered regions cannot be attributed to allelic differences. Additionally, 5 of these regions are potentially duplicated sequences with a sequencing depth exceeding 1.5-fold in the genome. Among the remaining 24 regions, 7 had an average sequencing depth of over 1.5-fold. After considering and excluding other factors, the methylation-based read clustering in the remaining 17 regions potentially indicates differences at the cellular level.

Furthermore, hifimeth-based CpG methylation calls can effectively capture differential methylation patterns across tandem repeats. For example, in the DXZ4 regions of CHM13, we observed distinct methylation patterns across the approximately 3kb repeat units in both the high-methylation group (HMG) and low-methylation group (LMG) reads (Figure 5B).

The high-accuracy hifimeth calls provide a means to uncover intricate single-molecule methylation patterns with high resolution, including alterations in methylation states within reads. While reads overlapping clustered bins were assigned to either the high-methylation group (HMG) or low-methylation group (LMG) based on the dominant clustered bins (weighted by bin CpG count), it is important to note that a read can exhibit different clustering patterns among the clustered bins. In HG002 and CHM13, we observed such methylation state switches on at least two reads in 1,010 and 4,623 distinct genomic regions, respectively (Supplementary Tables 8, 9). In the DXZ4 region of CHM13, we identified multiple clustered regions where the HMG reads displayed low methylation levels while the LMG reads exhibited high methylation levels. Three exemplary regions denoted by double-sided blue/red arrows (numbered 7, 8, and 9) in Figure 5B illustrate this phenomenon. Additionally, we observed an allelic differentially methylated CpG (chr7:135,168,118-135,168,119, p = 1.1E-4) in the CYREN gene in HG002 (Supplementary Figure 3), which displayed contrasting methylation levels compared to HMG and LMG group assigments. Furthermore, using the CYREN gene as an example, we observed a highly significant (p = 2.1E-11 by Fisher’s exact test) allelically differentially methylated CpG (chr7:135,169,124-135,169,125) at one of its splicing sites in HG002. Interestingly, the upstream and downstream CpGs did not exhibit significant allelic methylation level differences (Supplementary Figure 3). Similarly, in the CHM13 data, this isolated CpG demonstrated differential methylation between the HMG and LMG reads. Overall, these findings suggest that the regulation of CpG methylation can be intricate in native DNAs, and hifimeth calls have the capability to unravel methylation patterns with single CpG motif resolution.

## Discussion

The accuracy of CCS-based CpG methylation calling methods in recovering single-molecule methylation patterns has been limited. Our analyses have demonstrated that while these methods are effective in resolving the majority of CpGs in reads, their overall performance is hindered by low-confidence CpGs with less discernible signals. Excluding these low-confidence CpGs can improve the call accuracy to over 95%. However, this approach results in unresolved CpGs in reads and reduces the utilization rate of the data by over 29%. The reasons behind the signals of these low-confidence CpGs remain unclear, suggesting that optimization of sequencing techniques or incorporation of additional information may be necessary for further improvements.

Previous knowledge regarding the correlation of methylation frequencies among adjacent CpGs has aided in the quantification of site frequency. We have found that this correlation is likely rooted in the association of methylation states among CpGs on single molecules. Leveraging the strong association of methylation states within a short distance (<250 bp), our deep neural network model, hifimeth, has improved the accuracy of CCS-based 5mC detection to approximately 94.7%. This strategy holds theoretical promise for enhancing other single-molecule methylation calling technologies.

The high accuracy of hifimeth at the single-molecule level enables the study of intricate DNA methylation patterns. Our results demonstrate that adjacent genomic regions (even at distances of tens of base pairs) with similar methylation frequencies (unphased) can exhibit switched methylation levels at the single-molecule level. Remarkably, this can be detected without the need for phased variants. Furthermore, hifimeth calls can uncover individual CpG sites that are differentially methylated within a context of non-differential or contrary methylation, suggesting that DNA methylation can be regulated at the level of single motifs in native DNAs. Hifimeth has also successfully reproduced the complex methylation patterns (*26*) across the entire DXZ4 tandem repeats, which are challenging to resolve using NGS-based methods, with only approximately 25× CCS data.

The accuracy of CCS-based 5mC calling has been previously limited. However, by utilizing longer contexts that exploit the association of methylation states among adjacent CpGs, hifimeth has elevated the accuracy of CCS-based 5mC detection to a new level. Our analyses highlight that hifimeth calls can unveil sophisticated single-molecule methylation patterns, thereby advancing research in this field.

## Methods

### Reference genome and annotations

We obtained GRCh38.p13 (GCF_000001405.39 without alternative contigs) from NCBI RefSeq database as reference and the GENCODE v43 (*27*) (https://www.gencodegenes.org/human/) gene structure annotation for analyzing the HG002 data. For the CHM13 data, we downloaded the genome sequences and corresponding GENCODE v38 r2 annotation of T2T-CHM13 v2.0 (*28*) from https://github.com/marbl/CHM13. We aligned the zebrafish BS-seq data using bowtie2 v2.5.1 (*29*) to the closest zebrafish assembly (GCA_000002035.4) (*30*) from NCBI, achieving a mapping rate of approximately 50%. To improve the mapping rate, we performed de novo assembly of the CCS data using hifiasm v0.19.3-r572 (*31*), utilizing the primary contigs as a reference, resulting in a mapping rate of approximately 68.4% for the BS-seq data. We calculated the average CpG density, distances between immediately neighboring CpGs with no other CpGs in between (Figure 2B), and the cumulative ratio of immediately neighboring CpG pairs (Figure 2C) in the human genome using the whole-genome CpGs of the CHM13 v2.0 assembly. We simulated the Poisson distribution of immediately neighboring CpG distances in the human genome (Figure 2B) based on the average CpG density and the total CpG count in CHM13 v2.0. The average CpG density in zebrafish was determined using the hifiasm primary assembly mentioned above.

### BS-seq data analysis

We obtained the methylation frequency profiling results of ∼100× BS-seq data of HG002 analyzed by Bismark (*32*) from Oxford Nanopore Open Data Project (https://labs.epi2me.io/gm24385-5mc/). The zebrafish BS-seq was sequenced by our lab and also used in the study of ccsmeth (*8*). For the zebrafish BS-seq data, we employed Bismark v0.24.0 to perform whole-genome CpG methylation frequency profiling. We identified reference CpGs with a BS-seq read coverage ranging from ≥ 0.50 to ≤ 1.5-fold and methylation frequencies of 100% and 0% as fully methylated and fully unmethylated CpGs, respectively. This methodology was also applied to the nanopore data.

### Nanopore sequencing based methylation profiles

The nanopore methylation frequency profiling results of HG002 were obtained from Oxford Nanopore Open Data Project (https://labs.epi2me.io/gm24385-5mc/), utilizing Guppy v5.0.1 (Oxford Nanopore Technologies, Oxford, UK) and the dna_r9.4.1_450bps_modbases_5mc_hac configuration for base and methylation calling. Whole-genome methylation frequencies were profiled using modbam2bed (https://github.com/epi2me-labs/modbam2bed). For CHM13, we obtained the nanopore-based DNA methylation profiles as reported by Gershman et al. (*26*).

### Inputs and outputs of hifimeth single-molecule methylation calling module

Given a read *R= r*_0_*r*_1_ *… r_n_*_-1_ over the alphabet Σ = (A, C, G, T) and a CpG position *p* (0 ≤ *p* < *n*), we extracte *a k*-mer *K_p_ = r_p-h_r_p-h_*_+1_ *… r_p_*_+*h*-1_, where *h= k/*2. The missing bases are padded with *r_O_* (or *r_n_*_-1_) if *p< h* (or *n- p < h*). We constructe a graph *G_p_* for *K_p_*. This graph *G_p_* is the input of hifimeth. Every base *r_i_* in *K_p_* is represented as a node *N_i_* in *G_p_*. Every pair of adjacent bases *r_i_* and *r_i_*_+1_ is connected by a directed edge *e_i_ = r_i_* ⟵ *r_i_*_+1_. In this way *G_p_* contains *k* nodes and *k-*1 directed edges. The edges *e_i_* in *G_p_* have no features. Every node *N_i_* contains 15 features *t_i_* ∈ ℝ^15^, including base information, sequencing quality, forward strand IPD and PW, and reverse strand IPD and PW of *r_i_* respectively, summarized in a two-dimensional tensor of shape *k* x 15.

The output of hifimeth consists of two probabilities *p*_0_ and *p*_1_, indicating the probabilities of methylated and unmethylated states of this CpG site respectively. Cleary *p*_0_ *+ p*_1_ = 1. If a reference genome (optional) is given, the CCS bam, which includes kinetics information will be aligned to the reference by pbmm2 (*33*). Hifimeth will then output the corresponding positions of each called CpG on the reference and in the read, in addition to *p*_0_ and *p*_1_. Hifimeth also generates a BAM file that contains the 5mC base modification values in MM and ML tags, following the specifications outlined in the SAM tag specification document (https://samtools.github.io/hts-specs/SAMtags.pdf) (*34*).

### Design of graph convolutional deep neural networks

The deep neural networks of the CpG methylation calling module are depicted in Figure 2A. The module comprises 12 blocks and a total of 368,928 parameters. The first 11 blocks consist of a batch normalization layer (*35*), a graph convolution layer (*23*) and a rectified linear unit (ReLU) activation function. In the final block, the ReLU activation function is replaced by a global mean pool layer, followed by a feedforward layer. The batch normalization layers and the graph convolution layers are implemented using the pyg library (*36*).

### Training and testing dataset

The hifimeth training process involves using both native and treated datasets. The native training data were obtained by downsampling a flowcell of native human HG002 CCS reads with a library insert size of 15 kb (**ND**, ∼13.5×) from the study conducted by Baid et al. (*37*). Additionally, the treated NA12898 dataset from the study conducted by Ni et al. (*8*) was used for read model training. This dataset consisted of CpG-fully-methylated CCS data (FMD) derived from M.SssI-treated DNA and unmethylated data (UMD) obtained from PCR-amplified DNA. The testing CCS datasets include the training datasets augmented with five flowcells of native HG002 datasets from the study by Baid et al. (*37*) and the Human Reference Pangenome Consortium (HPRC), three flowcells of native CHM13 datasets with a library insert size of 20 kb, one flowcell of native zebrafish data (*8*), treated data of HG002 (training data for primrose), and M2&W2 (*13*).

### Training hifimeth

Hifimeth is trained using the cross-entropy loss function and the SGD optimizer with a learning rate of 0.1, weight decay of 10^(-5), momentum of 0.9, and Nesterov accelerated gradient. To expedite convergence during the training process, the learning rate is decayed by a factor of 0.1 after each epoch using the StepLR learning rate scheduler. Each hifimeth model undergoes training for two epochs. Optionally, a pretrained model can be used as a starting point for training.

### Context length and model performance test

To evaluate the effect of different k-mer lengths on model performance, we utilized a training set comprising 100,000 reads from the ND dataset. Positive and negative references were established by identifying fully-methylated and fully-unmethylated CpG sites using HG002 BS-seq data within the CCS reads. The models were assessed using 10,000 randomly sampled reads (∼1.5 Gb) from the ND dataset, excluding the 100,000 reads used for training. This evaluation was performed with three replicates to ensure robustness of the results.

### Training the K-20 and K-400 models

To train the K-20 model (20-mer context), we extracted positive and negative CpG features from 400 000 **FMD** and 400 000 **UMD** reads, respectively. The testing datasets consisted of 5 000 **FMD** and 5 000 **UMD** reads. The k-20 model served as a pretrained model for training the k-400 model, which was achieved by training hifimeth using a 400-mer context. To capture the CpG methylation correlation relationship in native DNAs, we constructed the training dataset for K-400 model primarily using native data, supplemented with a small portion of treated dataset. For the native data, we extracted 500 000 reads from the **ND** dataset and employed the K-20 model to call 5mC on these reads. Firstly, we selected CpG sites with methylation probabilities ≥ 0.9 or ≤ 0.1 from the K-20 model calls as positive and negative samples, respectively. Secondly, for the remaining CpG sites in these 500 000 reads, we included those that mapped to reference CpG sites with methylation frequencies ≥ 95% or ≤ 5% (identified by BS-seq) as positive or negative samples, respectively. Regarding the treated data, we extracted positive or negative samples from 60 000 reads of the **FMD** or **UMD** datasets, respectively, which constituted a third part of the training dataset. Additionally, we extracted CpG features from another 10 000, 5 000, and 5 000 reads from the **ND**, **FMD**, and **UMD** datasets, respectively, to serve as the testing dataset.

### Single-molecule CpG methylation correlation analysis

We randomly sampled 100 000 reads from one flowcell of native human data (**ND**) and another 100 000 reads from the flowcell of native zebrafish data. We applied the hifimeth K-20 model on the sampled reads, and only called the methylation states of the high-confidence CpGs in the reads, including CpGs with ≤0.1 methylation probability as negative and those ≥0.9 as positive. These high-confidence CpG methylation calls are estimated with ≥97% accuracy (Supplementary Figure 1A,B). The methylation correlation among high-confidence CpGs with different distances were calculated like the *r^2^*in linkage disequilibrium analysis. In this case, we treated the methylated and unmethylated states as two distinct alleles. We then calculated the r2 value between CpGs at a given distance using the following formula:

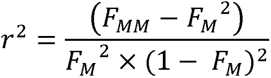

In the equation, *F_MM_* denotes the proportion of CpG pairs where both CpGs are methylated at the given distance. *F_M_* is the methylation frequency of the CpGs involved in CpG pairs at the given distance. To assess the statistical significance, we performed chi-squared tests and applied the Benjamini-Hochberg correction for multiple testing.

### Methylation frequency quantification module of hifimeth

Hifimeth incorporates two methylation frequency modes: read counting and site-frequency model modes. By default, hifimeth considers only reads with mapping qualities (MAPQ) equal to or greater than 10. In the read counting mode, the number of reads exceeding the positive probability threshold (0.5 by default) at a CpG is recorded as the methylated read count (*N_m_*), while the number of reads falling below the negative probability threshold (0.5 by default) is recorded as the unmethylated read count (*N_u_*). The methylation frequency at the CpG is then calculated as *N_m_ /* (*N_m_* + *N_u_*). The site-frequency model follows the same algorithm as ccsmeth (*8*). For training the frequency model, methylation frequencies obtained from hifimeth K-400 count mode on chromosomes 15 to 20, using ∼13.5× **ND** dataset, were used as the training set. Methylation frequencies on chromosomes 21 and 22 were used as the testing set.

### Assessment of single-molecule level performance

For the native datasets (HG002, CHM13, and zebrafish), the fully methylated and unmethylated CpGs identified by BS-seq (HG002 and zebrafish) or nanopore data (CHM13) were used as the reference positive and negative sites. The corresponding CpG sites identified in all mapped CCS reads with a mapping quality of ≥10 were used for assessment on each tested flowcell. For the treated datasets (NA12898, HG002, M2&W2), all CpGs from the sampled 10 000 **FMD** and 10 000 **UMD** reads were tagged as true positive and true negative, respectively. Three replicates of testing datasets were sampled for each of the three treated datasets using SAMtools v1.17 (*34*) and the ‘shuf’ command in bash. Methylation probability threshold(s) (default: 0.5) were used to distinguish positive and negative calls in hifimeth, ccsmeth v0.3.2, and primrose v1.3.0 (Pacific Biosciences, Menlo Park, USA). Based on the tagged CpGs in CCS reads, the methylation state calls were classified as true positive (TP), true negative (TN), false positive (FP), and false negative (FN). Then we calculated the AUC scores, precisions, accuracies, recall rates, and F1 scores for hifimeth, ccsmeth, and primrose calls on each tested dataset (native or treated) using Python scripts.

### Assessment with different probability thresholds

The entire dataset **ND** was used to assess the performance of ccsmeth, primrose, and the hifimeth K-20 model at the single-molecule level using probability thresholds from 0.01 to 0.99 (0.01 increment per step). The positive predictive value (PPV) and negative predictive value (NPV) at different thresholds were calculated as TP / (TP + FP) and TN / (TN + FN), respectively. When using two separate thresholds for positive and negative calls, any CpG with a methylation probability between the positive call threshold (>0.5) and the negative threshold (<0.5) was not assigned a methylation state. The data utilization ratio was calculated by dividing the number of called CpGs by the total number of CpGs in the analyzed tested reads.

### Assessment of methylation frequency quantification

To downsample the HG002 and CHM13 CCS data to desired sequencing depths (10-25×), we assessed the sequencing depth of each flowcell by using the ’samtools depth’ command on their mapped BAM files. We calculated the proportions of reads to be sampled for each desired depth and sampled an equally abundant number of bases per flowcell of each library type using the ’samtools view - s’ command. Using the downsampled datasets, we calculated the whole-genome methylation frequency profiles using the frequency quantification modules of hifimeth and ccsmeth for their respective methylation calls. For primrose calls, we used pb-CpG-tools v2.2.0. Both the read count mode and frequency model mode were applied to each software on each tested dataset. We then calculated the Pearson correlation coefficients between the methylation frequency profiles obtained by hifimeth, ccsmeth, and primrose and the corresponding BS-seq and nanopore-based results.

To assess phased methylation frequency profiling based on hifimeth, we merged the six flowcells of native HG002 CCS data and obtained its whole-genome phased variants (VCF file) from the study by Jarvis et al. (*38*). We performed down sampling at five different sequencing depths (10x-60x) as described above. For comparison, we obtained the mapped BAM file of ∼40x nanopore data of HG002 from the Oxford Nanopore Open Data project (https://labs.epi2me.io/gm24385-5mc-remora/). We used whathap v1.7 (*39*) to assign haplotype information (HP tag) to the reads in the aligned BAM files, including hifimeth methylation calls and nanopore methylation calls, using the phased VCF. CCS methylation frequency profiling was carried out using hifimeth with both count and frequency model modes for both paternal (HP:i:1) and maternal (HP:i:2) haplotypes. Phased methylation profiling was carried out using modbam2bed on the haplotype-tagged nanopore BAM file. Finally, correlation analyses were performed between CCS-hifimeth and nanopore results using CpGs with at least 5 reads per haplotype in both datasets for the paternal and maternal haplotypes separately.

### Allelic differential methylated regions

To detect differentially methylated regions (DMRs) between the two haplotypes, we used DSS v2.48.0 (*40*) with the CCS-hifimeth (count mode) and nanopore frequency quantification results. We applied thresholds of p < 0.001 and an allelic methylation difference greater than 0.2. The DMRs identified by both methods were then analyzed by taking the intersection using bedtools v2.30.0 (*41*). The DMRs were further classified as either maternal (-M) or paternal (-P) hypermethylated regions, based on which haplotype had a higher methylation frequency. To analyze their distribution across the genome, we divided the reference genome into 1 MB continuous non-overlapping windows. The proportions of maternal or paternal hypermethylated regions shown in Figure 5A were calculated by dividing the total length of maternal (or paternal) hypermethylated DMRs in each 1 MB window by 1 000 000..

### Methylation-based read clustering

To perform methylation-based read clustering, we followed a similar approach to the study by Gershman et al. (*26*) with some modifications. The following are the steps we took. We first split the reference genome into 200 bp continuous non-overlapping bins and screened those with high CpG density (≥5 CpGs per bin) for testing. The average methylation frequency (*M_bin_*) of each bin was calculated by dividing the total number of methylated CpGs by the total number of CpGs in all bin overlapping CCS read (MAPQ ≥ 10) fragments. In case that a read covers at least 5 CpGs in a bin, the methylation level of the read (*R_bin_*) was calculated by dividing the methylated CpG count by the covered CpG count. If the number of reads with *R_bin_*≥ *M_bin_* + 0.2 or *R_bin_*≤ *M_bin_*-0.2 accounted for ≥80% of the total reads covering the bin with ≥5 CpGs, reads with *R_bin_* ≥ *M_bin_* + 0.2 were assigned to the HMG and those with *R_bin_*≤*M_bin_* - 0.2 were assigned to the LMG. We further carried out Fisher’s exact test between HMG and LMG using their methylated and unmethylated CpG counts in reads. And bins with p*<*0.001 were identified as methylation clustered bins. When assigning the reads to HMG or LMG, the clustered bins were weighted using the number of CpGs they harbored. Reads with more HMG clustered bins were assigned to HMG, vise versa. We further screened clustered regions (≥ 400 bp) by merging adjacent bins with at most 200 bp distances using bedtools v2.30.0 (*41*). The circular plot shown in Figure 5A was drawn using Circos v0.69-9 (*42*). The methylation frequency plots and the plots showing the methylation status of CpGs on single molecules in Figure 5B and Supplementary Figure 3 were depicted by modbamtools v0.4.8 (*43*).

### Data availability

For HG002, two flowcells (20 kb library) of HG002 native CCS data can be obtained from the HPRC GitHub repository (https://github.com/human-pangenomics/HG002_Data_Freeze_v1.0), and the rest four flowcells (15 kb and 24 kb libraries) are available at https://console.cloud.google.com/storage/browser/brain-genomics-public/research/ /publication/sequencing (*37*). The treated HG002 data used for primrose model training was accessed from https://github.com/PacificBiosciences/primrose in March, 2023. The VCF file containing the whole-genome phased small variants of HG002 (*38*) is available at https://doi.org/10.18434/mds2-2578. The whole-genome CpG methylation frequency results (unphased) of HG002 by BS-seq and nanopore sequencing, together with the raw sequencing data are accessible from the Oxford Nanopore Open Data Project (https://labs.epi2me.io/gm24385-5mc). The mapped BAM file of ∼40× nanopore sequencing data are available at https://labs.epi2me.io/gm24385-5mc-remora. For CHM13, the three flowcells of CCS data (20 kb library) were obtained from the Telomere-to-telomere consortium CHM13 project (https://github.com/marbl/CHM13/tree/master) (*28*). The nanopore-based DNA methylation profiles for CHM13 (*26*) is available at https://zenodo.org/record/4739219. The flowcell of native zebrafish data and the treated NA12898 data can be obtained from https://ngdc.cncb.ac.cn/gsa using the Accession No. CRA010412 and HRA004180, respectively (*8*).

## Acknowledgment

We thank all those who generated and freely released the data analyzed in our present study. This study was funded in part by the National Key R&D Program of China (2022YFF1201900) and the National Natural Science Foundation of China (grant numbers 32270713, 62150048). We thank the Local Innovative and Research Teams Project of Guangdong Pearl River Talents Programme (2017BT01S138), CAMS Innovation Fund for Medical Sciences (2019-I2M-5-005) and Guangdong Basic and Applied Basic Research Foundation (2020B1515020057).

## Author contributions

C.L.X. conceived and designed this project. Y.C. and B.W. conceived, designed, and implemented the alignment algorithm. Y.C. integrated all the programs into the hifimeth pipeline and provided documentation. Y.Y.D., L.J.N, X.B. and Z.B.L. ran analyzed the performance of algorithms developed in this study. B.W., Y.C. and C.L.X. wrote the manuscript. All authors have read and approved the final version of this manuscript.

## COMPETING FINANCIAL INTERESTS

The authors have no competing financial interests to declare.

## Notes

### Competing Interest Statement

The authors have declared no competing interest.

