## Supplementary Figures for "High accuracy methylation identification tools on single molecular level for PacBio HiFi data"

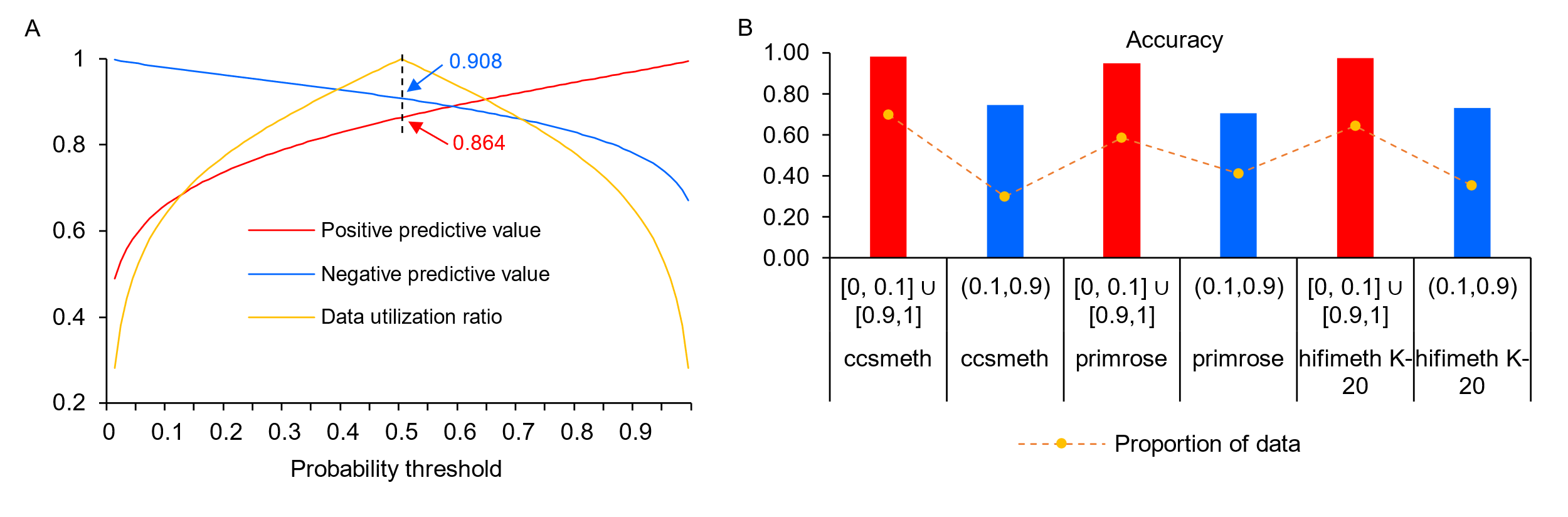

**Supplementary Figure 1. CCS-based methylation calling using different probability thresholds.** (**A**) Performance of hifimeth K-20 model using different software probability thresholds. (**B**) Accuracy and ratios of high-confidence and low-confidence CpGs. The same thresholds were used for different softwares: ≤0.1 for high-confidence negative calls and ≥0.9 for high-confidence positive calls; >0.1 and <0.9 for low-confidence calls.

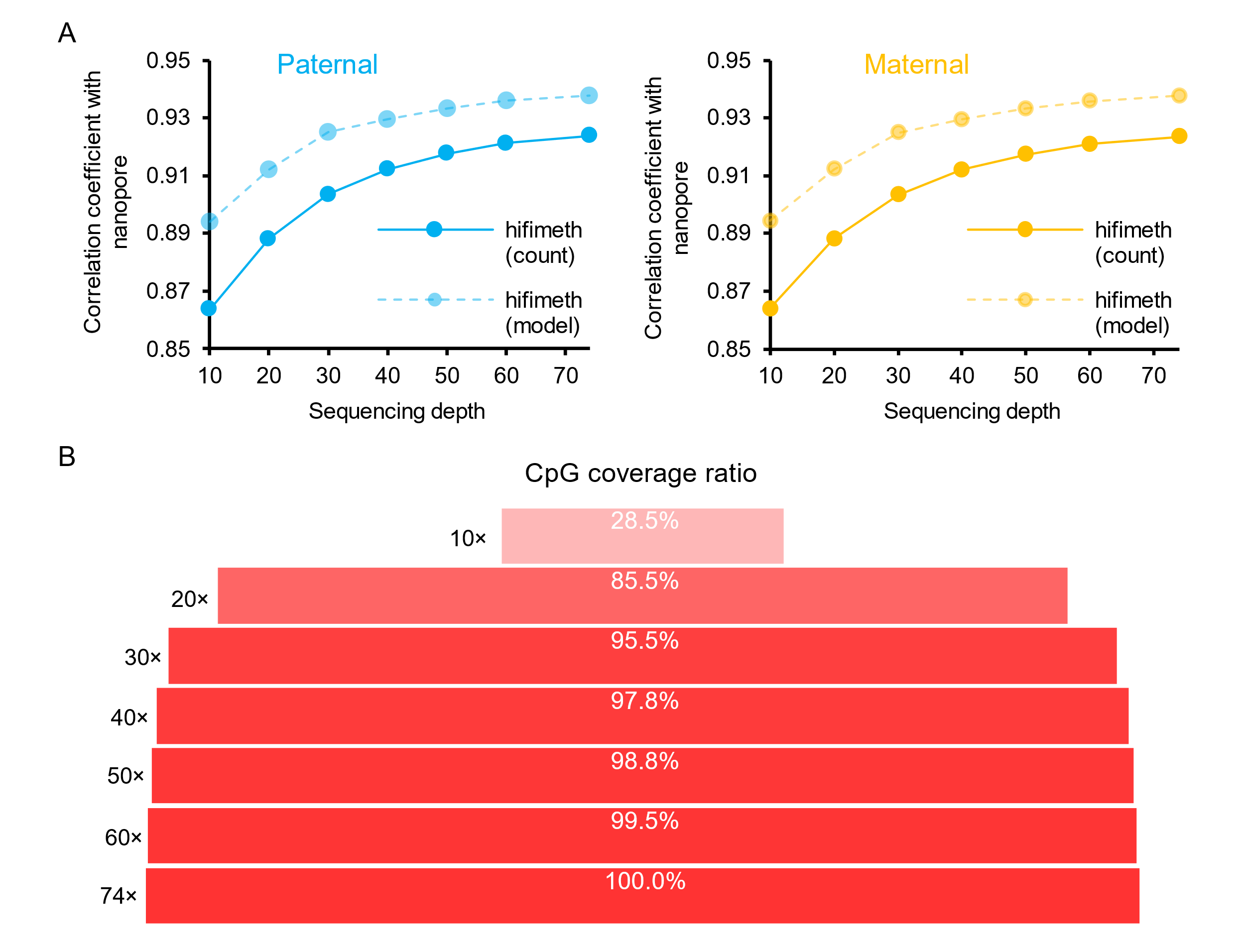

**Supplementary Figure 2. Phased methylation frequency assessment.** (**A**) Phased methylation frequency correlation between CCS datasets of different sequencing depth called with hifimeth and nanopore-based results. Three replicates of sampling were performed for 10-60 × depths and the average correlations are shown. (**B**) Proportions of CpGs quantified on both haplotypes at different CCS sequencing depths. Only CpGs with ≥ 5 called reads on either haplotype were counted as quantified. The proportions were calculated relative to the number of CpGs quantified at ~74 × depth. The correlation coefficients and number of quantified CpGs for each sampling replicate are in Supplementary Table 2.

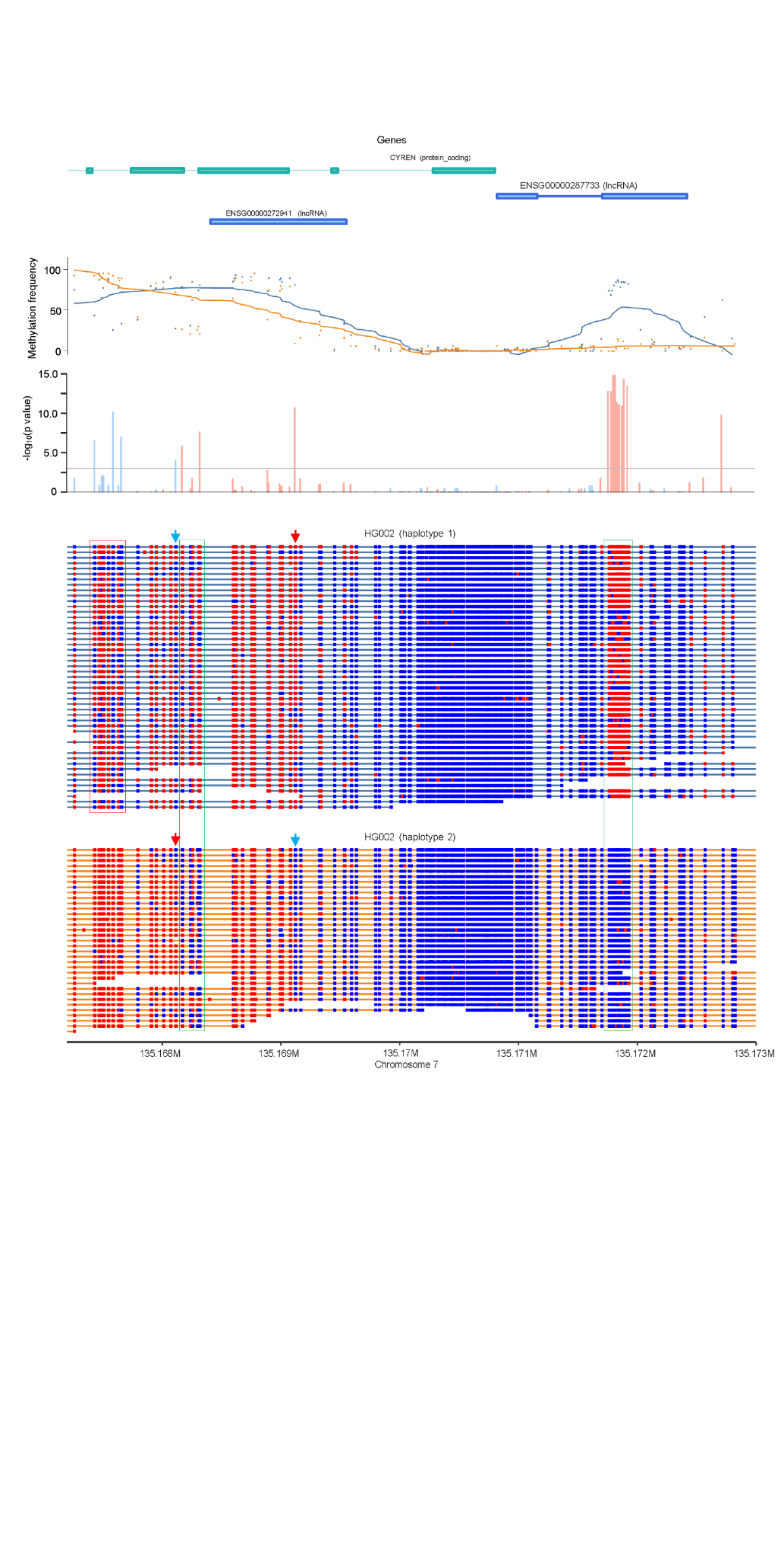

**Supplementary Figure 3. Phased methylation patterns at the *CYREN* gene in HG002.** From top to bottom: GenCode genet structural annotation, *CYREN* is on the reverse strand of the reference GRCh38 genome; methylation frequency panel: each dot denotes the paternal (blue, haplotype 1) or maternal (orange, haplotype 2) methylation frequency on the corresponding CpG, and the two curves show the smoothed average methylation frequencies calculated by modbamtools (Razaghi et al., 2022); histogram showing -log_10_(p value) of Fisher’s exact tests on the distribution of methylated and unmethylated calls on individual CpGs between the two haplotypes; the bottom panels show the single-molecule methylation patterns on reads assigned to paternal and maternal haplotypes, with blue and red bins showing unmethylated and methylated CpGs, respectively. In the bottom panel, the arrows denote two CpGs with significantly different methylation states compared to surrounding CpGs. Green frames indicate two clustered bins in the regions detected by methylation-based read clustering.
